## Supplementary figures and images for "Biochemical characterisation of human transglutaminase 4"

### Supplementary Figure 1

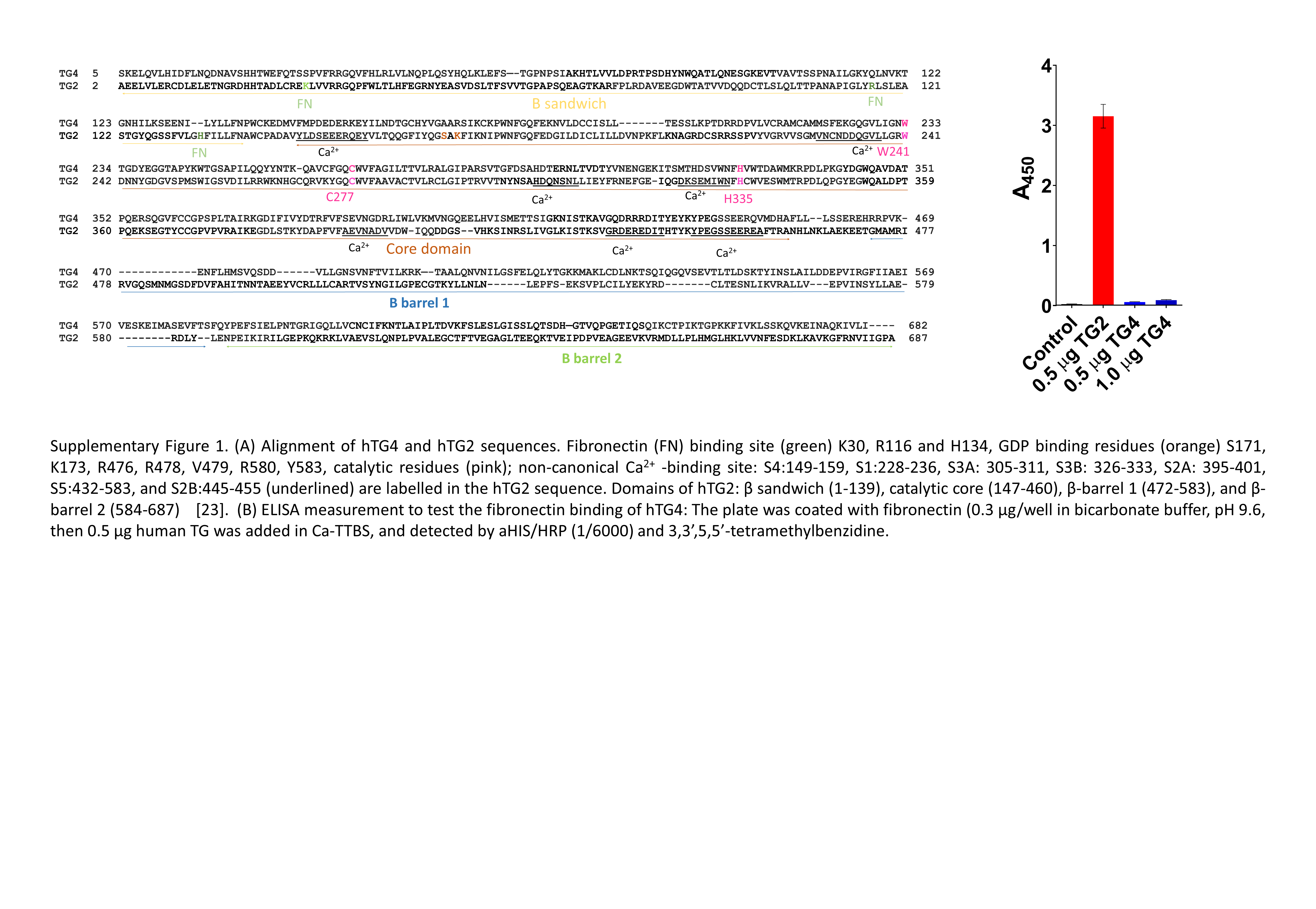
